# Skill-specific changes in cortical preparatory activity during motor learning

**DOI:** 10.1101/2020.01.30.919894

**Authors:** Xulu Sun, Daniel J. O’Shea, Matthew D. Golub, Eric M. Trautmann, Saurabh Vyas, Stephen I. Ryu, Krishna V. Shenoy

**Author notes:** Correspondence should be addressed to X.S. or K.V.S.

## Abstract

Animals have a remarkable capacity to learn new motor skills, but it remains an open question as to how learning changes neural population dynamics underlying movement^1^. Specifically, we asked whether changes in neural population dynamics relate purely to newly learned movements or if additional patterns are generated that facilitate learning without matching motor output. We trained rhesus monkeys to learn a curl force field^2^ task that elicited new arm-movement kinetics for some but not all reach directions^3,4^. We found that along certain neural dimensions, preparatory activity in motor cortex reassociated existing activity patterns with new movements. These systematic changes were observed only for learning-altered reaches. Surprisingly, we also found prominent shifts of preparatory activity along a nearly orthogonal neural dimension. These changes in preparatory activity were observed uniformly for all reaches including those unaltered by learning. This uniform shift during learning implies formation of new neural activity patterns, which was not observed in other short-term learning contexts^5–8^. During a washout period when the curl field was removed, movement kinetics gradually reverted, but the learning-induced uniform shift of preparatory activity persisted and a second, orthogonal uniform shift occurred. This persistent shift may retain a motor memory of the learned field^9–11^, consistent with faster relearning of the same curl field observed behaviorally and neurally. When multiple different curl fields were learned sequentially, we found distinct uniform shifts, each reflecting the identity of the field applied and potentially separating the associated motor memories^12,13^. The neural geometry of these shifts in preparatory activity could serve to organize skill-specific changes in movement production, facilitating the acquisition and retention of a broad motor repertoire.

## Introduction

Motor learning encompasses a wide range of phenomena, from relatively low-level mechanisms for calibrating movement parameters, to making high-level cognitive decisions about how to act in a novel environment^1^. Motor adaptation has been a long-standing and widely used paradigm for studying motor learning. Decades of behavioral studies have demonstrated many intriguing phenomena during motor adaptation, such as the error-driven calibration of movements, generalization of learned skills to a new context, savings (faster relearning) or memory retention, and interference between learning multiple skills^3,4,12,14–18^. Yet their neural mechanisms, in particular the underlying neural population dynamics, remain largely unknown.

In the field of motor control, neural population dynamics have provided foundational insight into activity patterns and computational principles not readily apparent at single-neuron resolution^19,20^. Recently, a dynamical system framework has started to help elucidate the neural basis of motor learning^5,8,21–23^. Collectively, these experiments have observed changes in neural population states related to the learning process. However, a remaining challenge is to dissociate neural population dynamics tightly linked to movement parameters (e.g., kinematics and kinetics) from other higher-level aspects of learning. For example, neural population correlates of motor memory (one consequence of learning) have yet to be identified. Through the lens of single-neuron activity, some studies found “memory” neurons — identified by persistent changes in their tuning properties following a washout period^9,24,25^—while other studies with similar but not identical tasks did not^26,27^. These results suggest that neural activity during learning can change in a way independent of movement parameters though what computational roles this type of changes could serve remains unclear.

Here we asked if all changes in neural population dynamics solely reflect the newly learned movements or, alternatively, if there are also dynamics that facilitate learning but are not tightly linked to motor output. To address this question, we designed a curl force field learning task that required generation of new movement kinetics for a subset of reaches while retaining the ability to generate original movements for other reaches (Figure 1a and see Methods).

**Figure 1.**
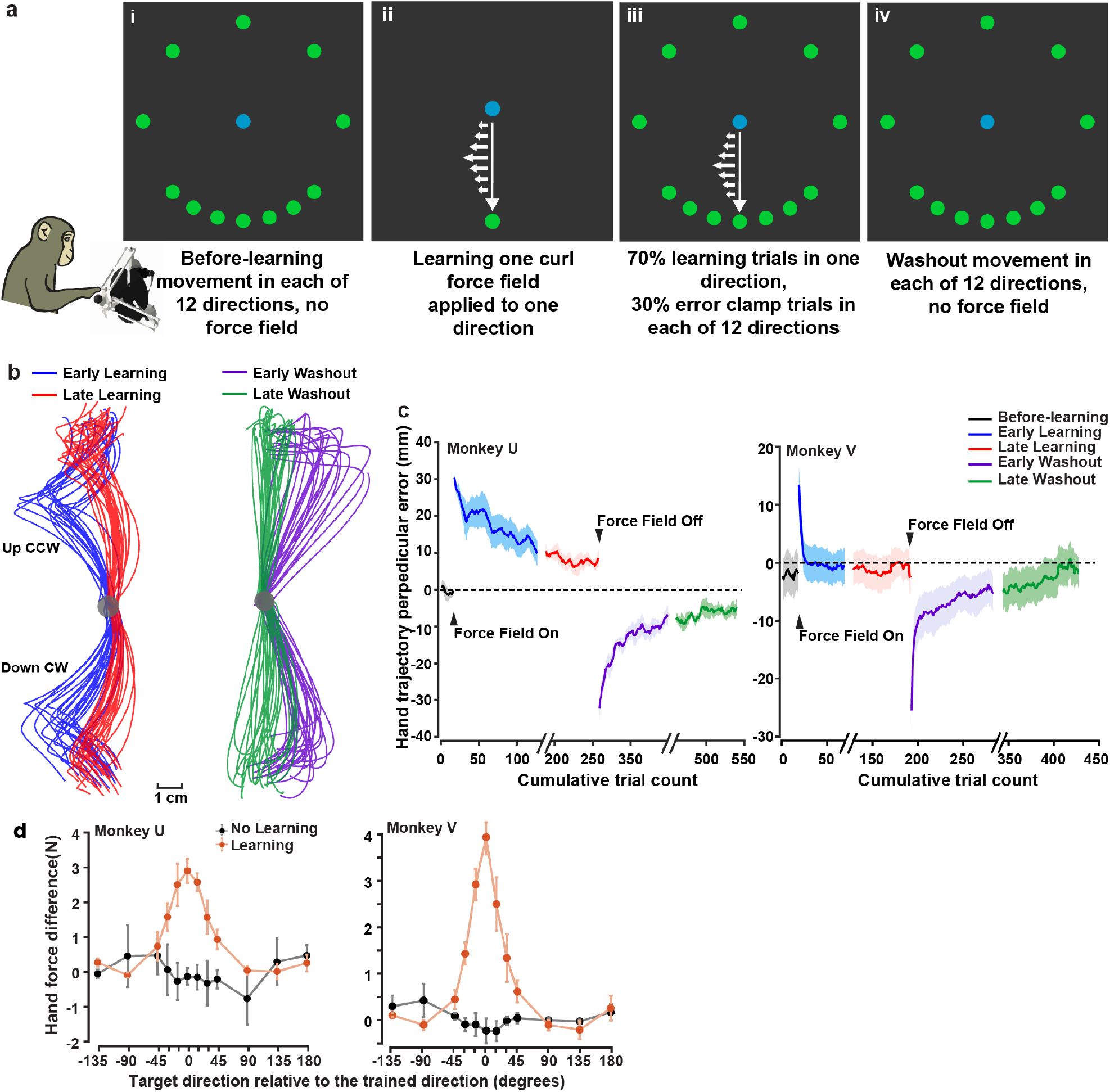
Task design and behavioral performance. **a**, Curl field learning and generalization task. Monkeys first performed delayed center-out reaches to each of 12 targets with no curl force field (block i: the before-learning block), followed by a learning block where one curl field was active during reaches to only one target (block ii; an example of a clockwise field applied to down reaches). After 150 successful learning trials (we found that 150 successes were sufficient for the decreasing hand trajectory error to plateau in each session), the task switched to an “error-clamp” block (see Methods) where 70% trials were the same as in block ii and 30% were randomly interleaved error-clamp trials for each of the same 12 reaching targets (block iii). The error-clamp paradigm constrained movements to a straight line toward the target and so held error feedback close to zero. It was used to measure changes in hand forces after-learning compared to the before-learning reaches as an indicator of the learning and generalization of new movement kinetics. The error clamp block was followed by a washout block in which the curl field was removed and monkeys needed to deadapt (block iv). Bottom-left inset, an illustration of the monkey controlling a haptic device. **b**, Example single-trial hand trajectories in one session applying a counterclockwise (CCW) force field to up reaches and another session applying a clockwise (CW) force field to down reaches. **c**, Behavioral indicators of learning and washout quantified by the hand trajectory deviation from a straight path (see Methods). Hand trajectory error was initially large and plateaued after decreasing (close to before-learning level) in late-learning and late-washout trials. Shaded area, s.e.m. across sessions. **d**, Behavioral generalization, measured by perpendicular hand force difference between error-clamp trials and before-learning trials (defined as the compensatory hand force), shows bell-shaped spatial pattern. Error bars, s.e.m. across sessions.

We trained two rhesus monkeys (U and V) to make delayed center-out reaches to each of 12 targets by controlling a haptic device (Figure 1a, block i), and then to learn a curl field applied during reaching to just one target (the trained target; Figure 1a, block ii). The force field perturbations were perpendicular to movement direction and proportional to hand speed. After monkeys had learned to reach straight again to the trained target in the force field (i.e. hand trajectory error plateaued after decreasing), reaches were again performed to each of 12 targets in “error-clamp” trials. In error-clamp trials, movement was constrained to a straight line toward the target, thereby clamping error feedback to zero (see Methods; Figure 1a, block iii). Error-clamp trials probed whether the newly-learned arm-movement kinetics, measured by the force applied to the haptic device, were transferred to nearby untrained targets to estimate the generalization of learning. Error-clamp trials were interleaved with learning trials (reaches to the trained target) to maintain the learned behavior. The error-clamp block was followed by a washout block in which the curl field was removed and the monkeys’ reaches exhibited aftereffects prior to deadaptation (Figure 1a, block iv). Monkeys displayed gradual behavioral learning and washout, quantified as a decrease in the perpendicular deviation of hand trajectory from a straight path (Figure 1b, c). The generalization of learning followed a bell-shaped spatial pattern: the amount of compensatory force falls off as the angle between the untrained target and the trained target increases (Figure 1d), consistent with human behavioral studies^3,4,14^.

We recorded neural activity in dorsal premotor (PMd) and primary motor (M1) cortex using Neuropixels probes (384 channels, Monkey V), Utah arrays (288 channels across three arrays, Monkey U), and V-probes (24 channels, Monkey V), which provided access to 100 - 300 neurons simultaneously (Neuropixels and arrays) or pooled over sessions (V-probes). We found that single-neuron activity during learning and washout were heterogeneous and complex (Supplementary Figure 1). This heterogeneity is consistent with previous reports^9,22,28^ and further motivated us to consider neural population activity and dynamics^19,20,29^.

### Reassociation-like changes of neural population activity in a 2D subspace

We first asked what signatures of curl field learning could be identified from the neural population activity, in particular, during movement preparation. Motor learning results in an update to the produced movement^1,30^, and preparatory neural activity (here −50 to +50 ms around the “go cue”) allows us to observe how the motor cortex may update movement preparation to generate the newly-learned behavior^8,23^. We used targeted dimensionality reduction (TDR^31^, see Methods) to estimate neural subspaces predictive of specific movement variables. Because precise control of hand forces is critical for learning curl field dynamics, we first focused on a low-dimensional subspace that captured across-trial neural variance related to horizontal and vertical hand forces. We constructed this subspace by regressing preparatory neural activity against initial hand forces (generated in the first 50 ms following movement initiation), because initial hand forces directly reflect the feedforward control of the prepared movement before sensory feedback arrives to motor cortex^32^. In this TDR subspace, before-learning neural states radially organized as a ring according to reach targets (Figure 2a) as expected^33,34^. During learning, preparatory states for the trained target rotated, from trial to trial, evolving towards the preparatory state of its adjacent target opposite to the curl field direction (Figure 2a, top-right inset). This rotatory shift towards the adjacent preparatory state is consistent with the motor system preparing to produce compensatory forces to counter the curl field; it resembles the “reaiming” strategy reported in a visuomotor rotation (VMR) task where monkeys’ motor preparatory activity rotates in the direction opposite the rotated visual feedback direction^6,8,35^. Importantly, we only used before-learning trials to build the TDR subspace. Hence, projecting population activity during and after learning (held-out trials) into this subspace yielded predictions of the new hand forces, which were strongly correlated with the real hand forces (Figure 2b) and exhibited high prediction accuracy (Figure 2c).

**Figure 2.**
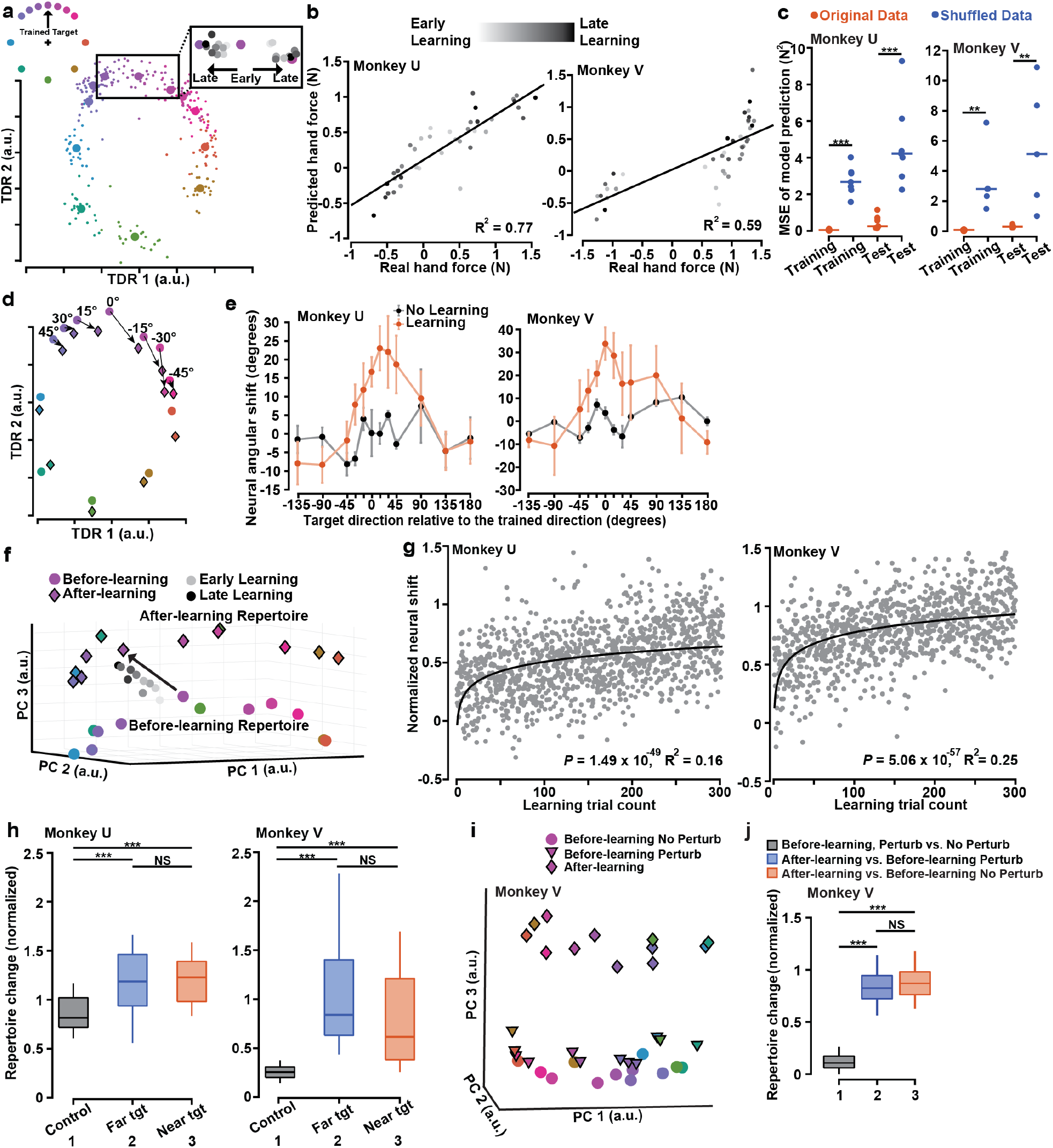
Patterns of preparatory activity in different neural state subspaces. **a**, In a 2D neural subspace constructed by TDR capturing the variance due to initial hand forces, preparatory states before-learning (color circles) radially organize as a ‘ring’ corresponding to reach targets (small circles: single-trial states; large circles: trial-averaged states). Top-left inset, color-coded reach targets when the trained target is up; Top-right inset, preparatory states during learning (grey and black circles) gradually rotating from the before-learning state of the trained target (middle) to that of its adjacent reaching target (left or right), in two example learning sessions (a CW or CCW force field applied to up reaches). **b**, Initial hand forces predicted by the 2D TDR preparatory states are correlated with real forces of the upcoming movement; the sign of hand force indicates its direction (Monkey U, *R*^2^ = 0.77 and *P* = 1.63 x 10^-13^; Monkey V, *R*^2^ = 0.59 and *P* = 1.92 x 10^-12^). Lighter dots, earlier learning trials; darker dots, later learning trials. **c**, Single-trial prediction MSE of initial hand forces is significantly smaller using original data than shuffled data (one-sided Wilcoxon rank-sum test: Monkey U, *P* = 0.0003 in both cases; Monkey V, *P* = 0.004 in both cases). For each monkey, left two columns, training-set (before-learning trials) prediction MSE; right two columns, held-out set (learning trials) prediction MSE. Control results (blue) are forces predicted by models built from training sets that have neural and behavior data shuffled. One datapoint per session. **d**, In the same TDR subspace, preparatory states of error-clamp reaches (color diamonds) consistently rotate in an opposite direction to the curl field direction for targets within 45 degrees from the trained target. **e**, Changes of preparatory states in the 2D TDR subspace reflect generalization of learning, quantified as the rotatory angles from before-learning to error-clamp neural states. Error bars, s.e.m. across sessions. **f**, Preparatory states of the trained target (grey circles) gradually shift away from the before-learning neural repertoire (color circles) and towards a new (significantly different) repertoire in error-clamp trials (color diamonds, the after-learning repertoire). The arrow points from the before-learning to the after-learning state of the trained target. Two-sided Wilcoxon rank-sum test: Monkey U, *P* = 3.22 x 10^-17^; Monkey V, *P* = 1.52 x 10^-13^. **g**, Single-trial neural shift during learning from the before-learning state along the axis that connects the centroids of before-learning and after-learning neural repertoires (see Methods). Solid line: linear-log regression (Monkey U, *R*^2^ = 0.16 and *P* = 1.49 x 10^-49^; Monkey V, *R*^2^ = 0.25 and *P* = 5.06 x 10^-57^). **h**, Preparatory neural repertoires change similarly for trained and untrained reaches. Black: control sessions in which monkeys did thousands of center-out reaches without any force field; blue (far tgt): far targets more than 45 degrees from the trained target in learning sessions; red (near tgt): near targets within 45 degrees from the trained target in learning sessions (one-sided Wilcoxon rank-sum test: Monkey U, *P*_12_ = 4.70 x 10^-5^, *P_13_* = 2.38 x 10^-8^, *P_23_* = 0.89; Monkey V, *P_12_* = 9.47 x 10^-9^, *P_13_* = 1.45 x 10^-7^, *P_23_* = 0.084). i, Preparatory states for before-learning no-perturbation (color circles), before-learning random-perturbation (color triangles), and after-learning (color diamonds) conditions projected to PCs 1-3. **j**, Larger preparatory repertoire changes occur to after-learning vs. before-learning noperturbation states (red), and after-learning vs. before-learning random-perturbation states (blue), than before-learning random-perturbation vs. no-perturbation states (black). *P_12_* = 1.83 x 10^-5^, *P_13_* = 1.83 x 10^-5^, *P_23_* = 0.37.

After learning, preparatory states for nearby, untrained targets also rotated towards their adjacent preparatory states that opposed the curl field (Figure 2d), and the amount of shift diminished with a similar bell-shaped spatial profile as seen for the behavioral generalization (Figure 1d, 2e). To our knowledge, this is the first direct evidence showing a neural population correlate of motor learning generalization, which has been predicted by previous work^24^. This reflection of behavioral generalization in neural preparatory states supports the theoretical framework that the internal model (representing a mapping from neural command to behavioral consequences) of reaching may be represented by a neural population code with a common set of basis functions shared by reaching to different targets^36,37^, and suggests that learning curl fields for one reach target modifies this shared basis, hence influencing untrained reaches in a spatially dependent manner.

This TDR analysis identifies a 2D neural subspace where shifts in preparatory states appear closely related to changes in behavior. This rotatory shift is consistent with the reassociation strategy observed during short-term brain-computer interface learning or the re-aiming observed during VMR learning, and suggests that reusing existing activity patterns may be a common strategy at least partially benefiting different learning contexts^6–8,38^.

### A uniform shift of neural population activity and the formation of new activity patterns

In contrast to a strict reassociation of existing activity patterns, we also identified prominent, unexpected neural changes along a nearly orthogonal dimension that did not directly parallel behavioral output. This was revealed by applying principal component analysis (PCA) to the neural population preparatory activity. We found that during curl field learning, preparatory states gradually shifted away from the before-learning repertoire (the set of neural states for reaching to all targets before learning), as visualized in the subspace of PCs 1-3 (Figure 2f grey circles) and quantified with full-dimensional neural data (Figure 2g, see Methods). The after-learning repertoire (the set of neural states for reaching to all targets in error-clamp trials) was significantly separated from the before-learning repertoire (Figure 2f). Strikingly, this shift of preparatory states was observed preceding reaches to both trained and untrained targets, and therefore was a “uniform shift” related to learning but not to the target specifically trained. This reveals components of preparatory activity that do not match the spatially-local behavioral generalization, in contrast to the rotatory changes in the 2D TDR subspace.

The presence of this uniform shift implies the emergence of new preparatory activity patterns (i.e., exploration of neural states unoccupied before-learning), which may constitute a neural repertoire change^7^. To test this hypothesis, we quantified the similarity between the after-learning and before-learning neural repertoires by measuring their normalized distance (see Methods). Consistent with the observed uniform shift (Figure 2f), we found that the preparatory repertoire changed substantially and uniformly for all reach directions (Figure 2h), which has not been discovered in other motor learning contexts^6–8,38^. Furthermore, we did not observe a uniform shift or repertoire change during VMR learning (Supplementary Figure 2). Thus, it appears that a distinct neural repertoire does not occur for all motor learning processes. During curl field learning, neural population activity shows a task-specific uniform shift, in addition to the reassociation strategy which seems to be shared by multiple learning contexts.

There are several trivial alternative explanations that could lead to the observed uniform shift and new neural repertoires yet bear no relation to a learning process. Here we discuss three sets of control experiments and analyses (see Methods) to rule out these potential reasons that are related to neural recording instabilities and behavioral changes irrelevant to learning. First, we compared repertoire change values in learning sessions to values in non-learning sessions when monkeys performed thousands of center-out reaches with the same task parameters but without applied force fields. The former was consistently and significantly greater than the latter (Figure 2h), which argues against within-session recording instabilities as the sole major contributor to the repertoire change, because otherwise similar repertoire changes should have been observed for both learning and non-learning sessions. Second, we did not find a preparatory neural repertoire change when we applied random pulse perturbation forces that simulated the magnitude of the curl field but did not involve learning (Figure 2i, j). This argues that the repertoire change did not result solely from preparing for upcoming movements that might be randomly perturbed by unpredictable forces. Third, muscle co-contraction could confound the result if it contributed to changes of neural activity not due to learning. Our electromyographic (EMG) recordings of muscle activation did not show obvious signs of co-contraction in late-learning and washout trials (Supplementary Figure 3), consistent with human behavioral studies^39^.

### The uniform shift reflects the identity of the applied curl field

So far, we have presented results from learning a single curl field. Next we asked whether different uniform shifts occur when learning different curl fields, and whether these shifts reflect the identity of learned fields. To address these questions, we trained monkeys U and V to learn different curl fields sequentially within the same session or over multiple sessions (see Methods). To track the activity of a neural population over multiple sessions, we selected a set of 71 neurons from Utah array recordings which exhibited highly similar waveforms across days^40^ (see Methods, Supplementary Figure 4a). We confirmed the stability of the selected units by examining their cross-day directional tuning (Supplementary Figure 4b). The stability of this neural population allowed us to directly compare the uniform shifts identified over multiple sessions.

To characterize uniform shifts when learning multiple force fields, we performed the following geometric analysis. For each learned curl field, we defined the “learning uniform-shift axis” as the vector pointing from the centroid of the before-learning neural repertoire to that of the after-learning repertoire. We then projected the before- and after-learning preparatory neural activity of each field onto its corresponding learning axis to visualize their geometric relationships. Last, we quantified the geometric relationships between uniform-shift axes by taking their dot products (see Methods).

We found that when monkeys (sequentially) learned two opposite curl fields applied to the same reach target, their preparatory neural states shifted in opposite directions, with respect to before-learning states (Figure 3a). Dot products of the uniform-shift axes were close to −1 and supported a nearly “antiparallel” relationship (Figure 3c green, Supplementary Figure 5a; see Methods). We also found that uniform-shift axes of clockwise or counterclockwise fields applied to different reach directions (up, right, or down) were close to orthogonal, in a pairwise fashion (Figure 3b). Dot products of uniform-shift axes in these cases were around 0 (Figure 3c purple). To further ground these curl-field-dependent uniform shifts quantitatively, we built a minimum distance decoder (see Methods), which predicted field types of held-out trials significantly better than by chance (Figure 3f). The existence of different uniform shifts for learning different curl fields and their geometric relationships provide evidence that the uniform shift is not an arbitrary change in neural population activity but relevant to learning.

**Figure 3.**
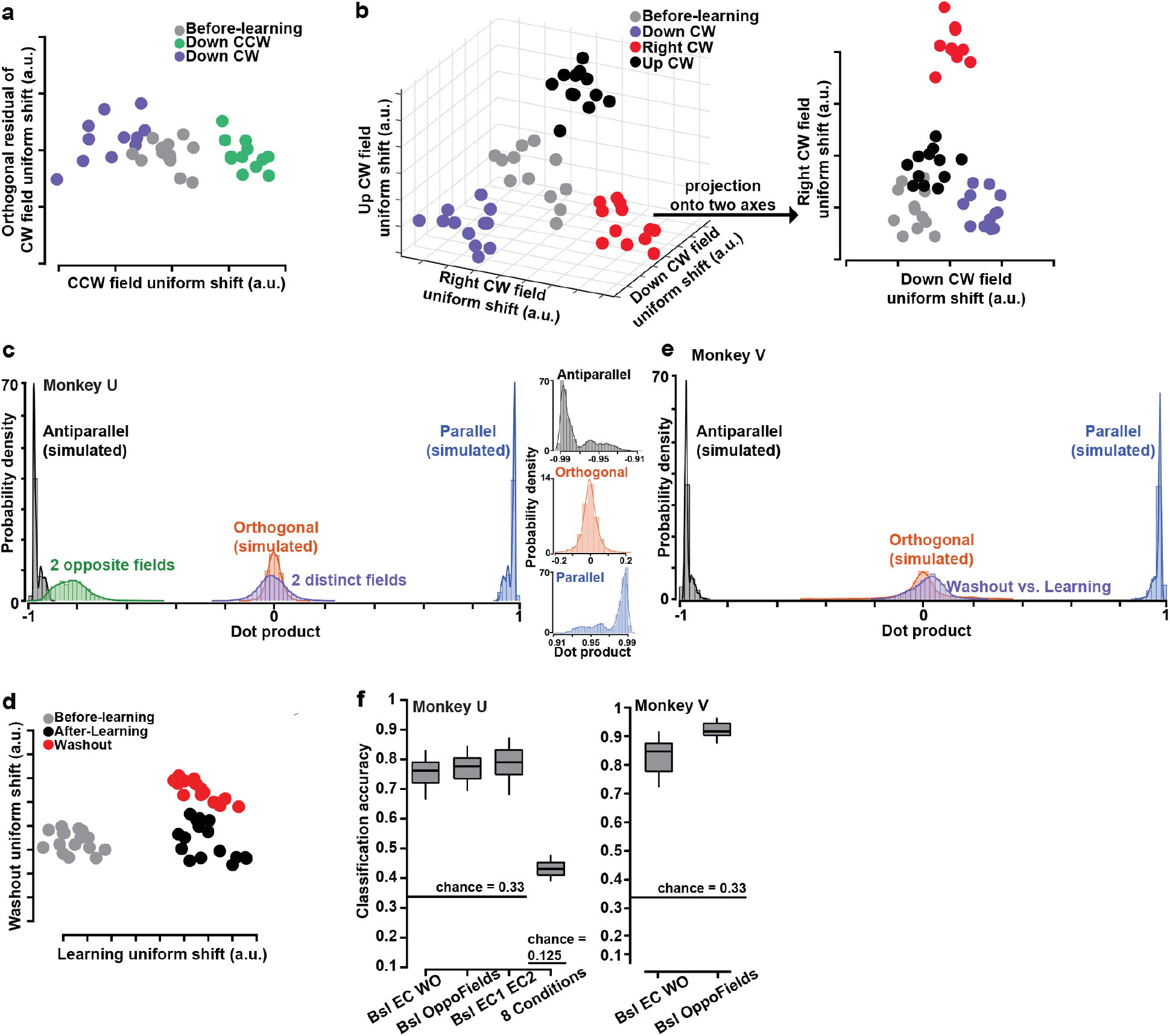
The uniform shift depends on and can predict curl field type. **a, b,** Observed geometric relationships of uniform shifts for learning different curl fields. **a**, Preparatory states projected to uniform-shift axes move in opposite directions when learning two opposite curl fields applied to one reach target. **b**, Uniform shifts for learning different curl fields applied to different reach targets are close to orthogonal. Left panel, preparatory states of learning three different fields projected onto three uniform-shift axes; right panel, projection of neural states in the left panel onto the uniform-shift axes of Down CW and Right CW fields. **c**, Distributions of dot products between uniform-shift axes when learning two opposite fields for one reach target (green) or two distinct fields for different reach targets (purple). We compare them with simulated distributions of dot products between uniform shifts predicted by “orthogonal” (red, around 0), “parallel” (blue, around 1), and “antiparallel” (black, around −1) relationships constructed from real data (see Methods). Inset, the zoom-in view of each hypothetical distribution. **d**, Observed geometric relationships of neural states along uniform-shift axes for learning and washout of one curl field. **e**, Distribution of dot products between uniform shifts when learning and washing out a curl field (purple), compared to hypothetical distributions predicted by orthogonal (red), parallel (blue), and antiparallel (black) uniform shifts constructed from real data. **f**, The accuracy for classifying before-learning vs. after-learning vs. washout trials (Bsl EC WO) and before-learning vs. curl field 1 vs. curl field 2 trials (Bsl OppoFields or Bsl EC1 EC2) from the uniform-shift component is significantly higher than by chance (0.333), using the minimum distance decoder and evaluated by cross validation (see Methods). One-sided signed rank test, *P* < 10^-50^ for all cases. The accuracy of classifying eight conditions (before-learning, learning five curl fields, and washout of two fields) from the uniform-shift component is significantly higher than by chance (0.125), evaluated by cross validation. One-sided signed rank test, *P* = 2.72 x 10^-51^.

### The uniform shift potentially retains a motor memory

Besides examining neural changes that accompany learning, the washout process provides a different lens to study the effect of learning, especially the retention of a motor memory^9,24,25^. We thus asked if any of the observed learning-related neural activity patterns persists after washout. After hundreds of washout trials, both monkeys reverted to before-learning arm-movement behavior (Figure 1b, c, Supplementary Figure 3, Supplementary Figure 6a). Correspondingly, preparatory states in the 2D TDR subspace gradually rotated back towards the before-learning states (Figure 4a, b). Along the learning uniform-shift axis, by contrast, preparatory states remained separated from the before-learning repertoire (Figure 4c, d). Interestingly, we identified a distinct uniform shift occurring during washout that was almost orthogonal to the learning uniform-shift axis (Figure 3d, e, Supplementary Figure 5b, with hypothetical models in Supplementary Figure 7e). This suggests that washout is not simply the reverse of learning but instead washout states shift in a new direction. From the learning and washout uniform shifts, a minimum distance decoder could predict before-learning, learning, and washout conditions significantly better than by chance (Figure 3f). These results suggest that the uniform shift of preparatory activity could potentially retain a motor memory that persists after washout.

**Figure 4.**
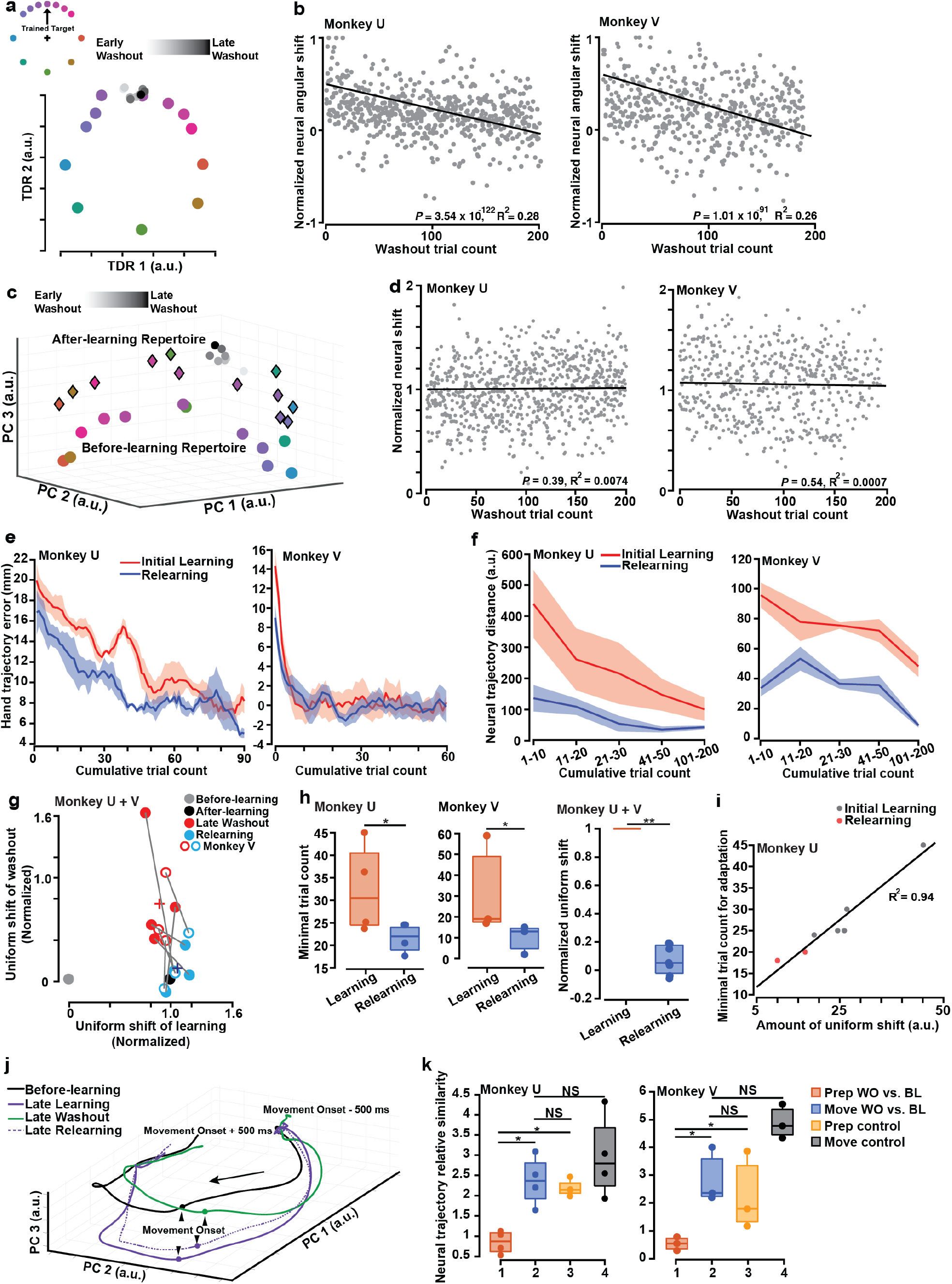
The correlation between the uniform shift and motor memory retention. **a**, In the same TDR subspace as in Figure 2a, preparatory neural states (grey and black circles) gradually rotate from the after-learning state back towards the before-learning state during washout. Inset, color-coded reach targets when the trained target is up. **b**, The angular differences between washout states and the before-learning state gradually decrease on a single-trial basis (solid line: linear regression. Monkey U, 3.54 x 10^-122^; Monkey V, *P* = 1.01 x 10^-91^). **c**, Preparatory washout states (gray and black circles) remain away from the before-learning repertoire (color circles) and close to the after-learning repertoire (color diamonds) along the uniform-shift axis. **d**, The distance between washout states and the before-learning state along the learning uniform-shift axis does not show significant trend of increase or decrease (solid line: linear regression. Monkey U, *P* = 0.39; Monkey V, *P* = 0.54). Each dot is a single trial. Normalized against the uniform shift of learning (see Methods). e, Hand trajectory errors are smaller during relearning than during initial learning (one-sided rank-sum test: Monkey U, *P* = 2.30 x 10^-5^; Monkey V, *P* = 0.003). f, Distance between learning neural trajectories and the late-learning trajectory decreases over trials (red) but is larger than distance between relearning neural trajectories and the late-relearning trajectory (blue). One-sided rank-sum test: Monkey U, *P* = 3.43 x 10^-4^; Monkey V, *P* = 8.23 x 10^-5^. **e, f**, Shaded area, s.e.m. across sessions. **g**, Centroids (circles) of late-washout and relearning states projected onto the learning and washout uniform-shift axes (7 sessions from 2 monkeys), normalized against the uniform shift of learning in each session. Late-washout states are significantly different from the learning state (one-sided signed rank test, *P* = 0.039 for x < 1 and *P* = 0.0078 for y > 0). Relearning states are significantly different from late-washout states (one-sided rank-sum test, *P* = 0.0055 for Δx > 0 and *P* = 0.0012 for Δy < 0), but not significantly different from the learning state (two-sided signed rank test, *P* = 0.22 for x compared to 1 and *P* = 0.47 for y compared to 0). Crosses, the means. **h**, The minimal trial count until compensatory hand forces reach 80% of the mean late-learning forces is significantly smaller during relearning than initial learning (one-sided rank-sum test: Monkey U, *P* = 0.043; Monkey V, *P* = 0.050). Correspondingly, the uniform shift from the late-washout state to the relearning state is significantly smaller than from the beforelearning state to the learning state, along the learning uniform-shift axis (normalized against the uniform shift of initial learning in each session; one-sided signed rank test, *P* = 0.0078, 7 sessions from 2 monkeys). **i**, The non-normalized uniform shift during learning or relearning is correlated with learning rate (71 neurons tracked over 5 sessions, including 2 relearning sessions). **j**, Neural trajectories of before-learning, late-learning, late-washout, and late-relearning conditions (500 ms before movement onset to 500 ms after movement onset). The late-washout trajectory (green) is farther from the before-learning trajectory (black) during preparatory than movement epoch. k, The similarity between late-washout and before-learning neural trajectories is significantly higher during movement (blue) than preparatory (red) period (one-sided rank-sum test: Monkey U, *P_12_* = 0.029; Monkey V, *P_12_* = 0.050), and the former can compare to the similarity between before-learning (yellow and black) neural trajectories (twosided rank-sum test: Monkey U, *P_23_* = 0.49, *P_24_* = 0.34; Monkey V, *P_23_* = 0.40, *P_24_* = 0.10). BL: before-learning. WO: late-washout. Prep: preparatory. Move: during movement.

To further elucidate the role of preparatory activity in motor memory retention, we next asked if the same after-learning neural states could be revisited when relearning the same field (see Methods). Monkeys U and V relearned the curl field faster with smaller perpendicular hand deviation error compared to the initial learning (Figure 4e). Neural trajectories during relearning approached the well-learned trajectory in fewer trials than during initial learning (Figure 4f). During relearning, preparatory neural states moved from the washout repertoire to the relearning repertoire that was not distinguishable from the learning repertoire (Figure 4g). Not only were the same after-learning neural states revisited within each relearning session, we also found that uniform shifts of the same curl field in two sessions 18 days apart were close to parallel (Supplementary Figure 5c green). This underscores that a specific uniform shift of preparatory states is associated with each field.

We also explored if the amount of uniform shift during learning and relearning (i.e. the distance between learning/relearning states and the before-learning state along the learning uniform-shift axis, see methods) is connected to behavioral learning or relearning rate. Associated with faster behavioral relearning, the amount of uniform shift was significantly smaller during relearning (Figure 4h). Quantification of uniform shifts in five sessions showed that a smaller shift was correlated with faster learning (Figure 4i, 71 neurons tracked across five sessions, see Methods). Together, our findings provide evidence for the contribution of the uniform shift to motor memory retention that could benefit learning.

### The uniform shift is a feature of preparatory neural activity

To determine whether the uniform shift is a unique feature of preparatory activity, we applied PCA to examine changes of peri-movement states (0 to 100 ms following movement initiation) after learning or washout. After learning, while peri-movement states also shifted away from before-learning ones (Supplementary Figure 8a, b), the shift was local and matched the bell-shaped behavioral generalization (Supplementary Figure 8a, c). Correspondingly, the peri-movement neural repertoire change was also local (Supplementary Figure 8d). After washout, peri-movement states reverted to before-learning patterns (Supplementary Figure 8e, f), directly reflecting the deadapted motor output. Consistent with this distinction between preparatory and peri-movement states after washout, the similarity between late-washout and before-learning neural trajectories (measured as the distance between washout and after-learning neural trajectories relative to the distance between washout and before-learning trajectories, see Methods) was lower during movement preparation than execution (Figure 4j, k). The uniform shift appears to be a preparatory phenomenon and disappears once movement is initiated.

## Discussion

In this study, we identified motor cortical activity patterns in different neural population dimensions that reflect distinct components of learning new arm movements. In a 2D TDR subspace, we found reassociation-like changes of preparatory neural states that seem to be shared by multiple learning contexts^6–8^. We also discovered a surprising uniform shift that may be specific to the context of learning curl force fields.

The occurrence of uniform shifts provides evidence for formation of new activity patterns during short-term motor learning. Conventionally, the circuit structure or connectivity of an existing network has been thought to constrain the patterns that its neurons are capable of exhibiting, which may limit its capacity for short-term learning^41–43^. Here our results suggest that the motor system may be more flexible than previously thought, and can generate novel activity patterns during short-term learning in order to quickly adapt to a changing environment^38^. To adapt to the new force environment when learning a curl force field, subjects need to acquire new movement kinetics (Supplementary Figure 6b). This task demand is a major feature differentiating curl field learning from other short-term learning contexts in which reassociation of existing neural activity patterns is sufficient to support behavioral learning^6–8^. The demand for learning new movement kinetics may lead the motor system to engage new neural activity patterns.

Furthermore, the uniform shift of preparatory neural states occurs to reach directions where the movement is unaltered during learning, which indicates that uniform shifts do not directly lead to motor output and so the resulting new activity patterns live in the “output-null” subspace of preparatory activity^44^. If we make the approximations that the observed preparatory activity is mainly generated in PMd and that a major output of PMd is M1^45^, the uniform shift is consistent with neural changes found previously in the “M1-null” subspace of PMd for rapid learning^22^. This output-null property of the uniform shift is also corroborated by our results that peri-movement states do not shift uniformly but change only for the altered movements. Assuming that at least part of the peri-movement neural dynamics generates behavioral output^19,20,29^, the lack of a uniform shift during movement execution further supports the idea that it does not directly lead to movement output.

Although the observed uniform shifts do not directly match behavioral output, our results show that they reflect the identity of learned curl fields and may facilitate the separation of corresponding motor memories that could otherwise interfere with each other. The behavioral phenomenon of interference has been widely studied using curl force field perturbations^4,16^. Interference occurs when opposing force fields that alternate or switch randomly from trial to trial are applied. Consequently, neither force field is learned. Recent behavioral studies have found that when people plan for, or imagine, different movements associated with different curl fields, they can learn multiple skills without interference that would otherwise hamper learning^12,46^. Here we propose that the uniform shift separating preparatory neural states may comprise a neural mechanism for this reduction of behavioral interference. That is, motor cortical preparatory activity sets the initial state of a dynamical system, whose subsequent evolution generates movement activity following a certain neural trajectory^19,29,34,47–50^. When subjects make the same reaching movement in different curl fields that switch randomly, if modifications due to learning are made around a single neural trajectory initiated from the same preparatory neural state, interference may occur. Instead, introducing a uniform shift to the preparatory activity separates initial states for seeding the local neural dynamics that would evolve in those regions of state space to produce distinct movement kinetics^51^. We speculate that uniform shifts could thus reduce interference between these neural dynamics and enable the generation of the appropriate motor signals.

Altogether, we propose a mechanism involving the uniform shift for motor memory separation, retention, and faster relearning. First, though the uniform shift is not tightly linked to motor output, it takes neural population activity to a state that might transition to the after-learning state more easily than from the before-learning state. Second, even after washout, the motor system preserves the uniform shift to retain a short-term memory, which could facilitate faster relearning. We observed distinct uniform shifts when learning different curl fields and they can separate these motor memories to potentially reduce interference and achieve more precise movement control. These roles of the uniform shift substantiate the idea that neural population activity has certain dimensions that organize internal dynamics for computation and pattern generation, but do not resemble the final behavioral output^44,52,53^. In contrast to the mechanisms proposed here, it is possible that changes we observed reflect learning-induced changes in a different brain region, e.g., cerebellum or basal ganglia. In future work, it may be possible to directly and specifically perturb the uniform shift by precisely manipulating the neural population activity (i.e., modulate specific neural population dimensions), which could provide a causal test of this hypothesized mechanism^54–56^.

## Methods

All surgical and animal care procedures were performed in accordance with National Institutes of Health guidelines and were approved by the Stanford University Institutional Animal Care and Use Committee.

### The single-trial temporal structure and the curl force field configuration

Two adult male rhesus macaques *(Macaca mulatta)* U (14 kg, 8 years old) and V (10 kg, 8 years old) were trained on the curl force field learning and generalization task. A diagram of the task design is shown in Figure 1a. Monkeys griped the handle of a haptic device (delta.3 haptic device, Force Dimension, Switzerland) with their right hands to control the movement of a cursor in a twodimensional plane displayed on the screen in front of them. Monkeys performed point-to-point delayed reaching task using the haptic device: they initiated each trial by holding their hand with a center target for 450 ms. Then a second target 120 mm (105 mm for Monkey U) away from the center showed up on the screen which served as the endpoint of the movement the monkeys were asked to make. This target originally vibrated in place for a variable delay period (200 – 650 ms, uniformly distributed) and stopped vibrating as a “go cue” which instructed the monkeys to reach. In curl field trials, the haptic device was programmed to produce forces on the monkey hands as they performed the point-to-point reaching movement. The magnitude and direction of the force depended on the velocity of hand movement according to Equation (1)

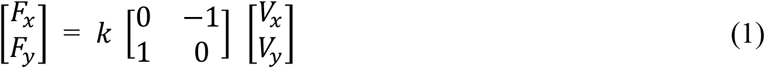

where *k* was set equal to ±14 N m^-1^ s for Monkey V and ±12 N m^-1^ s for Monkey U. The sign of *k* determines the direction of the curl force field: positive *k* for counterclockwise (CCW) fields and negative *k* for clockwise (CW) fields.

We applied either CW or CCW curl field to three reaching directions in this study: up, down, and right (i.e. six different fields). In one session, only one reaching direction had curl field active. Hand position and velocity were measured at 1k Hz by the haptic device. Hand forces were measured at 30k Hz by a load cell mounted to the haptic device and then down-sampled to 1k Hz during behavior data processing.

### Block design of the principal task

Throughout the study, a session means a day’s worth of experiments, spanning several hours. In every learning session, monkeys first made delayed reaches from the center of the workspace to one of 12 peripheral targets without force field in a before-learning block (block i, 20 trials per reaching target). Then the learning block (block ii) started where a curl field was active for one reaching target (the trained target). After 150 successful learning trials, monkeys showed behavioral adaptation (Figure 1c) and the task entered the error-clamp block (block iii) where 70% trials were the same as in the learning block and 30% were randomly interleaved error-clamp trials^57^ (20 trials per reaching target). The error-clamp trials were used to probe any change in hand forces without causing visual error feedback because the hand trajectory was ensured to be straight. After the error-clamp block (except for multiple-field-learning experiments, see below) was a washout block (block iv) very similar to the before-learning block in which there was no force field.

Throughout block iii, to assess feed-forward learning of the curl field, the error clamp was rendered by the haptic device when monkeys reached to each of the same 12 targets as in block i (Figure 1a). In these trials, the hand movement was confined to a simulated mechanical channel with a spring constant (stiffness) of 10,000 N/m and damping constant of 150 Ns/m^4,58^.

In learning and washout blocks, in order to encourage the monkeys to learn or unlearn the curl field (rather than accept making highly curved and inefficient movements to targets), we introduced a path efficiency check, where we automatically failed a trial if the hand trajectory deviation (perpendicular to the straight-line trajectory to the reach target) exceeded a bound. In all trials throughout all sessions, monkeys were required to finish each reach movement within 700 ms or the trial would be counted as a failure.

### Relearning experiment

To investigate the neural substrate of motor memory retention (the behavioral indicator of which is defined as savings), monkeys were exposed to the same curl field for a second time post-washout (Monkey U, four sessions; Monkey V, three Neuropixels sessions). In these sessions, monkeys did at least 500 washout trials before relearning to make sure the learned behavior was washed out as completely as possible.

### Double-field-learning experiment

To identify and investigate neural components of learning and generalizing a specific type of curl field, in three sessions, Monkey U learned two opposite curl fields sequentially for the same reaching target (up, right, or down); in one Neuropixels session, Monkey V learned two opposite curl fields for the same reaching target (up). Monkeys were exposed to each force field with a learning block and an error-clamp block, similar to a typical single-field-learning session. No washout trials were performed between the two fields.

### Center-out reach control experiment

This control experiment was conducted to verify that the changes of neural population activity patterns were learning-related, not merely due to the instability of recording. In control sessions (Monkey U, n = 3; Monkey V, n = 2), Monkeys U and V made thousands of delayed center-out reaches to one of the 12 targets without any curl force field. The task had the same temporal duration as in a learning session. For data analysis, we used trials for which the trial IDs matched those in learning sessions to measure neuronal changes due to temporal drift.

### Random-perturbation control experiment

This control experiment was conducted to verify that the changes of neural population activity patterns were learning-related, not merely because of generating larger muscle forces in a new environment. In one Neuropixels session, Monkey V first performed center-out reaches to one of the 12 targets without any perturbation force (the before-learning, no-perturbation block). Then in 50% of all center-out reach trials, the haptic device applied a pulse perturbation force, either to the left or to the right of the reaching direction, which simulated the force magnitude of the curl field but the perturbation direction was not predictable (the before-learning, random-perturbation block). So Monkey V could not learn to prepare for the pulse perturbation but instead needed to generate compensatory forces after sensing it. In this block, trials with and without perturbation forces were randomly interleaved. All successful trials from the before-learning, no-perturbation block, and perturbation trials in the random-perturbation block were used in data analysis.

### Visuomotor rotation (VMR) experiment

The VMR experiment has been described previously^8^. Two male adult Monkeys, R (15 kg, 12 years old) and J (16 kg, 15 years old), were trained to perform a delayed center-out reach task to one of eight targets. A VMR perturbation was introduced to all eight reach targets during the learning block. Each monkey had two 96-electrode Utah arrays, one implanted in PMd and one in M1. The arrays were implanted five years and seven years for R and J respectively prior to the experiments. In control sessions for the VMR experiment (three sessions per monkey), Monkeys R and J made thousands of delayed center-out reaches to one of the eight targets without any VMR.

### Neural data collection

In a standard sterile surgery, Monkey V was implanted with a head restraint and a recording cylinder (19 mm diameter) located over M1 and caudal PMd (coordinates A16, L15) on the left hemisphere. Three Utah arrays (1 mm-long electrodes, spaced 400 μm apart, Blackrock Microsystems) were implanted for Monkey U eight months prior to the experiments. Extracellular spikes were recorded using three 96-channel Utah arrays implanted in PMd, lateral M1 and medial M1 (Monkey U), 24-channel Plexon V-probes (Monkey V, specifically: PLX-VP-24-15ED-100-SE-100-25(640)-CT-500), and 384-channel Neuropixels Phase 3A silicon probes^59^ (Monkey V). Monkey U datasets included 10 recording sessions (four sessions of the relearning experiment, three sessions of the double-field experiment, and three sessions of the center-out control experiment); Monkey V datasets included seven Neuropixels recording sessions (three sessions of the relearning experiment, one session of the double-field experiment, two sessions of the center-out control experiment, and one session of the random-perturbation experiment) and two sets of V-probe recordings for two different curl fields (20 sessions in total).

### Neural data pre-processing

For Utah array recordings, voltage signals were band-pass filtered from each electrode (250 Hz – 7.5 KHz). These signals were processed to detect “threshold crossing” spikes. We detected spikes whenever the voltage crossed below a threshold of −3.5 times the root-mean-square voltage. For V-probe and Utah-array recordings, spike sorting was performed offline using a custom software package^44^ (available online as MKsort; https://github.com/ripple-neuro/mksort/); stable single units and multi-unit isolations were included. Around 250 units from 20 V-probe recording sessions were sorted by MKsort and used in the analysis. Around 300 single- and multi-units from the three Utah arrays passed the sorting criteria in each session. For Neuropixels recordings, the original data were automatically spike sorted with the Kilosort spike sorting software and then manually curated with the ‘phy’ gui^60^ (https://github.com/kwikteam/phy). Around 1000 units from seven Neuropixels recording sessions (100 – 200 units per session) were sorted by KiloSort and ‘phy’ and used in the analysis. For V-probe recordings, neurons recorded over multiple sessions were pooled together for the same curl field because these sessions shared a common task structure and configuration, which resulted in at least 100 units per curl field. Despite the various spike-detection or sorting methods, we acquired consistent analysis results.

### EMG data collection and pre-processing

For Monkey U, surface EMG recordings (Trigno EMG Systems, Delsys Inc.) were made from triceps brachii, biceps brachii, and posterior deltoid. For Monkey V, surface EMG recordings were made from the trapezius, lateral deltoid, pectoralis, biceps brachii, extensor carpi radialis, and flexor carpi radialis. EMG signal was processed by the signal envelopes taking the upper and lower peaks smoothed over 80-sample intervals.

### Data analysis

The behavioral, EMG, and neural data were analyzed offline using MATLAB 2017b and 2019a (MathWorks). All analyses pooled together PMd and M1 recordings, and used the full-dimensional neural data unless otherwise specified in the following sections.

### Behavior quantification

To examine behavioral learning, generalization, and washout, two measures were used: 1) kinematic error (the lateral hand trajectory deviation), and 2) the kinetic change (hand force difference). The kinematic error was calculated on each adaptation movement as the maximum perpendicular error (MPE) of the hand path relative to a straight line joining the center and the target locations of the movement. For both monkeys, the MPE for all blocks in each session was rolling average over 10 trials to reduce the single-trial noise while preserving the temporal resolution of learning, with the sign flipped appropriately so that errors from CW and CCW field trials could be appropriately combined. The mean and s.e.m. of MPE were then computed across all sessions (Figure 1c).The kinetic change was calculated as the perpendicular hand force difference between error-clamp trials and their corresponding before-learning trials for the same reaching target (averaged over 100 – 200 ms after movement onset around which time the perpendicular hand force reached the peak). Note that throughout the study, ‘perpendicular’ means ‘perpendicular to the reach direction’.

### Principal component analysis (PCA)

To perform PCA on neural data, we constructed the data matrix ***R*** of size *N_neuron_* × (*N_condition_ · T*), with *N_neuron_* the number of neurons, *N_condition_* the number of conditions (defined by reaching directions and force field type) and *T* the number of time points per condition during preparatory epoch (−50 to +50 ms from go cue for all neural state analyses except that for neural repertoire analyses it was −400 to −100 ms from movement onset), peri-movement epoch (0 to 100 ms from movement onset for all neural state analyses except that for neural repertoire analyses it was 0 to 600 ms from movement onset), or a whole trial (−500 to 500 ms from movement onset): time points were spaced 1 ms apart for neural trajectory analyses or averaged over a 100 ms bin to generate the neural states within a certain time window; the response of a given neuron was centered by subtracting its mean response from the firing rate at each time point; the centered neural data were then averaged across all trials for each condition. The condition-averaged neural data matrix was passed to the SVD function to compute the principal components (PCs). The movement onset for each reach was determined via a speed threshold of 35 mm/s for Monkey V and 40 mm/s for Monkey U.

### Targeted dimensionality reduction (TDR)

We applied the TDR approach^31^ to identify lowdimensional subspaces capturing variance related to the behavioral variables of interest. To construct the TDR space, we used multivariable, linear regression to determine how various behavioral variables affect the responses of each unit. We first centered the responses of a given unit by subtracting the mean response from the firing rate at each time point. The mean was computed by combining the unit’s responses across all trials and times. We then described the centered responses of neuron i as a linear combination of several behavioral variables:

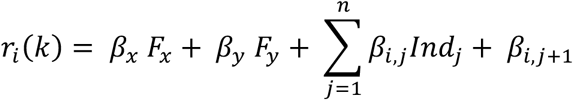

where *r_i_* is the centered, trial-averaged response of unit *i* binned over a certain time window (same as in the PCA procedure) on condition *k. F_x_* and *F_y_* are the horizontal and vertical hand forces on condition *k. Ind_j_* is a binary indicator of the trial type: it is 1 if condition *k* is the error clamp trial for curl field *j* and 0 otherwise. The last regression coefficient *β_i,j+1_* captures variance independent of the listed behavioral variables. The 2D hand-force subspace was built by regressing full-dimensional preparatory neural states against hand forces averaged over 50 ms after movement onset (before sensory feedback signals arrived at motor cortex) without binary indicators (i.e. *n* = 0). We also built a 3D TDR model that incorporated an indicator of the trial condition (before-learning vs. after-learning error clamp trials) as an additional regressor (i.e. *n* = 1), and this model revealed a uniform shift during learning along the third dimension similar to the PCA results (data not shown).

To estimate the regression coefficients, we constructed the following matrix ***M*** of size *N_condition_ × N_Coeffi_*, where *N_coeffi_* is the number of regression coefficients to be estimated, and *N_condition_* is the number of conditions for which the neural data were recorded. The first *N_Coeffi_* −1 columns of ***M*** each contain the condition-by-condition values of one of the behavioral/task variables (the regressors). The last column consists only of ones to estimate ***β_j+1_***. Given the *N_condition_ × N_neuron_* matrix of neural firing rates for all the conditions, ***R***, the regression model can be written as:

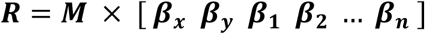

And the regression coefficients can be estimated as:

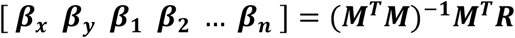

We then projected the neural data into the regression subspace by multiplying the pseudoinverse of the ***β*** coefficient matrix with the neural data matrix ***R***.

### Measurement of changes to the neural repertoire

We applied the approach proposed previously^7^ to top 10 PCs of the neural population data. To briefly summarize, we quantified the neural repertoire change as the distances between each after-learning neural activity pattern in the error clamp block and its nearest neighbors among all of the before-learning patterns, normalized by the variance of the before-learning repertoire. Values near zero indicated repertoire preservation and larger values indicated repertoire change. We measured and compared the neural repertoire changes for curl field learning sessions (using before-learning and error-clamp trials), center-out control experiment sessions (using center-out reach trials matching the trial IDs in a learning session), and random-perturbation experiment sessions (using before-learning no-perturbation trials, before-learning random-perturbation trials, and after-learning error-clamp trials).

### Definition of uniform-shift axes and quantification of uniform neural state shift

In the visualization of preparatory neural states projected to the first three PCs (Figure 2f), we observed that all the after-learning states were separated from the before-learning states (a uniform shift). We thus defined uniform-shift axes by the following steps. We took the centroids of preparatory states (i.e. *N_neuron_* × 1 vectors without PCA) of all reach directions in the before-learning block, after-learning error-clamp block, and late-washout block. The axis that connected before-learning and after-learning centroids was defined as the “learning uniform-shift axis”; the axis that connected after-learning and washout centroids was defined as the “washout uniform-shift axis”. We orthogonalized these axes against the two TDR axes where we found rotatory neural shifts (see Main). The statistical test of Figure 2f was performed on trial-averaged neural states of before- and after-learning conditions for all reach targets projected onto the learning uniform-shift axis. In Figures 3a, b, d, and 4g, we orthogonalized uniform-shift axes before projecting neural activity onto them. In Figures 2g, 4d, and 4i, we projected full-dimensional neural population activity in learning, washout, and relearning trials of the trained target onto the learning uniform-shift axis to quantify their distance from the before-learning state along this axis. Figure 4g shows centroids of the trial-averaged, late-washout and late-relearning neural states projected onto the learning and washout uniform-shift axes (7 sessions, Monkeys U and V). Normalization was performed against the distance between before- and after-learning centroids in each session (defined as the “uniform shift of learning”, see Figure 4d, g) unless otherwise specified.

### Measurement of geometric relationships between uniform-shift axes

We took the dot products between different uniform-shift axes as defined above to measure their geometric relationships: values close to 1 indicated that the two axes were close to parallel, −1 indicated an antiparallel relationship, and 0 indicated orthogonality. This geometric analysis could hypothetically reveal at least four different relationships between uniform shifts when learning distinct curl fields: (1) “parallel overlapping” happens when uniform shifts for two fields are in the same direction (their dot product equals 1), and after-learning neural repertoires are mixed (Supplementary Figure 7a); (2) “parallel non-overlapping” results from uniform shifts in the same direction (their dot product equals 1) with after-learning neural repertoires segregated for different fields (Supplementary Figure 7b); (3) “antiparallel” describes uniform shifts in opposite directions (their dot product equals −1, Supplementary Figure 7c); (4) “orthogonal” represents uniform shifts in independent directions (their dot product equals 0, Supplementary Figure 7d). We bootstrapped trials of each condition and repeated the measurement of dot products between the bootstrapped uniform-shift axes to estimate distributions of dot products. The control distributions of dot products for parallel, antiparallel, and orthogonal axes were constructed by the following procedure: 1) Parallel/antiparallel case: for each pair of before-learning and after-learning centroids, we measured the dot products between uniform-shift axes using bootstrapped trials and the uniform-shift axis using all trials, because without noise intrinsic to the data, they should be truly parallel. The antiparallel distributions were constructed from taking the inverse of the parallel distributions but with a separate, independent set of resampling. 2) Orthogonal case: we orthogonalized the trial-averaged uniform-shift axes for learning two curl fields applied to two different reach directions (or axes for learning and washing out one curl field), denoted as 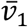 and 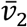. We also denoted their corresponding bootstrapped, orthogonalized uniform-shift axes as 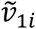 and 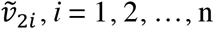. We then measured the dot products between 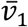 and 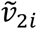, as well as between 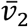 and 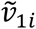, because without noise intrinsic to the data, they should be truly orthogonal.

### Minimum distance decoder

The minimum distance decoder used half of all trials as training trials to find the centroids of preparatory neural states as defined above, and decoded the condition type (before-learning, after-learning, or washout for a given curl field) and the curl field type based on to which centroid the neural state vector of the test trial was closest (i.e. smallest Euclidean distance). The decoding performance was evaluated by cross validation.

### Measurement of minimal neural trajectory distance (Figure 4f)

To estimate the minimum neural distance between different conditions over time, we performed a modified Euclidean distance analysis^49^. We selected points on one of the two neural trajectories we were comparing (learning: relearning trajectory; before-learning: learning trajectory; washout: relearning trajectory) and calculated the Euclidean distance between that point and every point on the second trajectory, in the first 10 PCs. We elected to use 10 PCs that account for over 90% of the variance of the data in all data sets. We selected the minimum Euclidean distance across all points on the second trajectory as our estimate of neural distance between the two trajectories at that time. This ensured that we would never overestimate the distance between the trajectories due to misalignment in time. A low distance indicated that the second trajectory was more similar to the first trajectory than when the distance was high.

### Measurement of relative neural trajectory similarity (Figure 4k)

The relative neural trajectory similarity was a metric to compare whether the washout neural trajectory was more similar to the before-learning or the after-learning trajectory. It was quantified as the ratio of neural trajectory distance (see the last section) between washout and after-learning trajectories over that between washout and before-learning trajectories averaged over a certain time window: for preparatory period, the time window was −50 to +50 ms from go cue; for movement period, it was 100 to 200 ms after movement onset. A larger value indicates higher similarity to the before-learning neural trajectory. Control analyses measured this metric using the before-learning neural data of which trials were split into two random halves, one half serving as the before-learning trials and the other serving as the sham washout trials.

### Tracking neurons over multiple sessions

To examine the relationship between uniform shifts of learning multiple force fields, we selected neurons that showed up in the same Utah array channel over five successive days. We evaluated the cross-day similarity of waveforms for each sorted neuron by 1) binning the waveform data into a 40D vector for each sorted neuron per session), and 2) calculating the Pearson correlation coefficient between waveforms on day 1 vs. day 2, day 1 vs. day 3, and so on^40^. The null correlation coefficient was generated from comparisons between waveforms that were known to be from different neurons (i.e. neurons from separate channels). Neurons with cross-day waveform correlation significantly higher than the null correlation were selected (i.e. higher than 95% quantile of the null correlation). At last, we confirmed the stability of the selected neurons by visualizing their waveforms and examining their directional tuning PSTHs in before-learning trials over days. Waveforms and PSTHs of example neurons over multiple sessions are shown in Supplementary Figure 4.

### Minimum trial count for learning or relearning (Figure 4h, i)

It was defined as the trial count during learning or relearning when maximum compensatory hand force within 100 ms after movement onset first achieved 80% of the same compensatory force averaged over the last 50 successful learning trials (i.e. the late-learning trials).

### Statistics

To test the single-trial learning and washout of neural states, we fitted regression curves of neural shifts against the trial count (Figures 2g, 4b, and 4d). To test the trend of gradual learning and gradual washout of neural states binned over trials, we applied the Cuzick’s test (Supplementary Figure 8b, f). It is Cuzick’s extension of the one-sided Wilcoxon rank-sum test to assess trend in data with three or more ordinal groups. Because we did not assume that the data followed a normal distribution, we applied the Wilcoxon rank-sum test tos compare groups of data and signed rank test to compare a group with a null mean value, using the one-sided test where appropriate. We used bootstrap to repetitively subsample the original data to generate before- and after-learning neural states for acquiring control distributions for the dot products of uniform-shift axes (details described in a former section). For all tests, we used *P* = 0.05 as the significance threshold. *P* values are included in each figure legend. Error bars are due to quantification from multiple trials per condition, from multiple sessions, or from multiple reach directions × multiple sessions.

## Author contributions

X.S. conceived the project and designed the experiments with input from D.J.O. and M.D.G. X.S. conducted the V-probe, Utah array, and EMG recordings. X.S. and D.J.O. conducted the Neuropixels recordings, with extensive assistance from E.M.T. S.V. collected the VMR data. X.S. performed data analysis with significant assistance from D.J.O. and M.D.G., as well as input from S.V. S.I.R. led the recording chamber and array implantation surgeries. X.S. wrote the manuscript with input and editing from all authors. K.V.S. was involved with all aspects of the research.

## Acknowledgements

We thank the members of the Shenoy lab at Stanford University for comments and discussions on the methods and results, and T. Fisher for help with experiments and comments on analyses. We thank M. Risch, M. Wechsler, and R. Reeder for expert veterinary care. We thank B. Davis for administrative support. We thank W. L. Gore Inc. for donating Preclude artificial dura used as part of the chronic electrode array implantation procedure. X.S. was supported by a Stanford Interdisciplinary Graduate Fellowship, a Stanford Bio-X Honorary Fellowship, and Stanford Department of Biology Funding. D.J.O was supported by a US National Science Foundation graduate research fellowship and a Stanford Graduate Fellowship. M.D.G was supported by an NIH K99/R00 award NIMH-K99MH121533. E.M.T was supported by an NIH NRSA grant 1F31NS089376-01, a Stanford Graduate Fellowship, and an NSF IGERT grant 0734683. S.V. was supported by an NIH F31 Ruth L. Kirschstein National Research Service Award 1F31NS103409-01, an NSF Graduate Research Fellowship, and a Ric Weiland Stanford Graduate Fellowship. K.V.S. was supported by the following awards: NIH Director’s Pioneer Award 8DP1HD075623, Defense Advanced Research Projects Agency (DARPA) Biological Technology Office (BTO) ‘‘NeuroFAST’’ award W911NF-14-2-0013, the Simons Foundation Collaboration on the Global Brain awards 543045, the Office of Naval Research N000141812158, and the Howard Hughes Medical Institute.

## Data availability

The data that support the findings of this study are available from the corresponding author upon reasonable request.

## Code availability

The code for the repertoire change analysis is available on github (https://github.com/mattgolub/bci_learning). All other code is available from the corresponding author upon reasonable request.

---

**Supplementary Figure 1.**
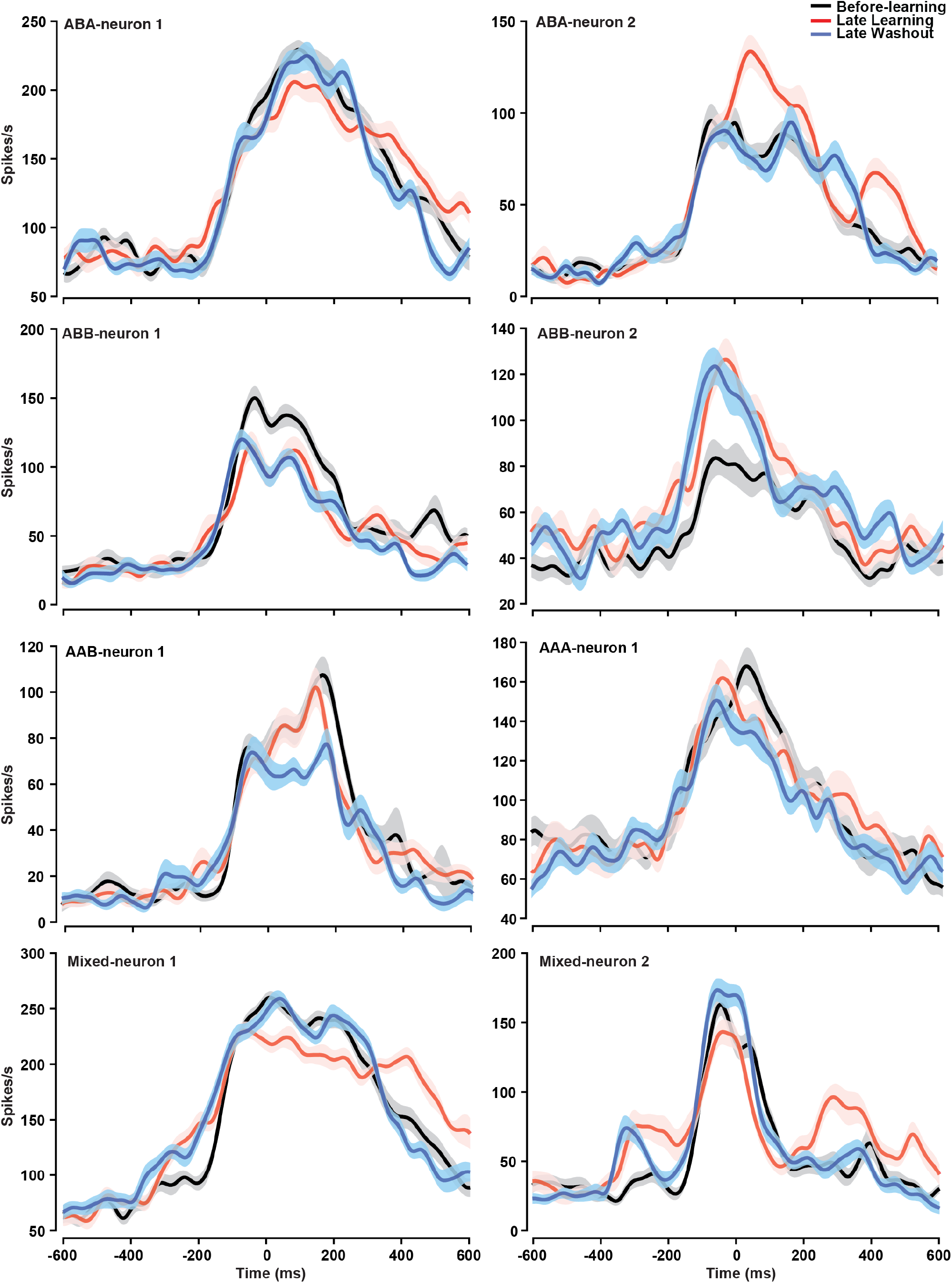
Example single neuron peristimulus time histograms (PSTHs). Comparing single-neuron activity across before-learning (black), learning (red), and washout (blue) blocks, we found neurons that changed activity during learning and reverted to before-learning activity after washout (ABA neuron); some neurons changed activity during learning and remained changes after washout (ABB neuron); some did not change activity until the washout block (AAB neuron); some maintained the same across all blocks (AAA neuron); neurons also exhibited mixed patterns of changes during preparatory and peri-movement epochs (e.g., mixed-neurons 1 and 2 show ABB pattern during preparatory epoch but ABA pattern during peri-movement epoch). This heterogeneity of activity was consistent with classic observations^9,22,28^. Shaded area, s.e.m across trials.

**Supplementary Figure 2.**
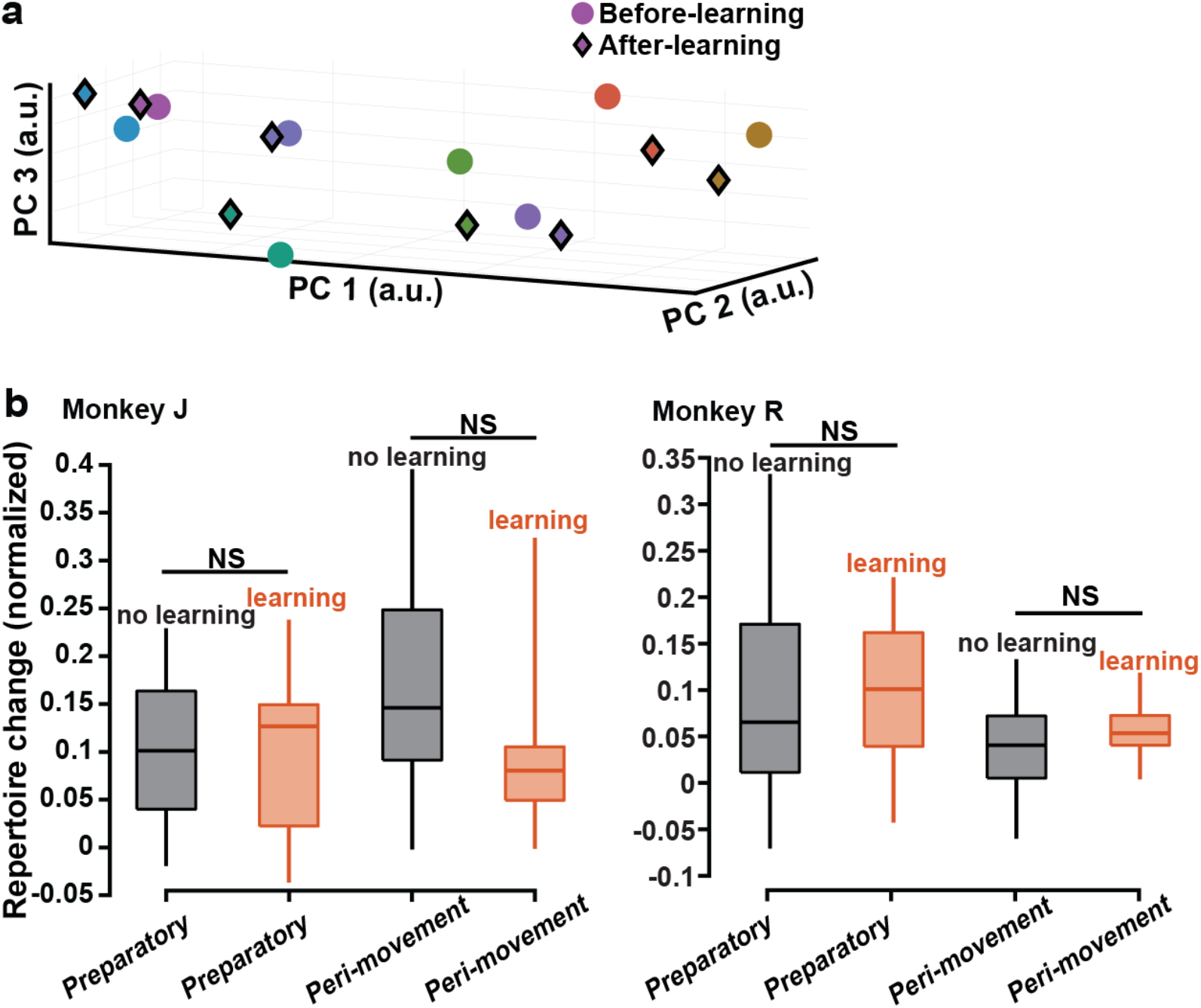
VMR control results. **a**, Preparatory neural states for VMR learning projected to PCs 1-3. After-learning states (color diamonds) are mixed with before-learning states (color circles). One example session. **b**, Preparatory and peri-movement neural activity patterns do not show repertoire change during VMR learning. One-sided Wilcoxon rank-sum test, *P* > 0.1 for all comparisons; three learning sessions (red) and three control sessions (black) for both monkeys.

**Supplementary Figure 3.**
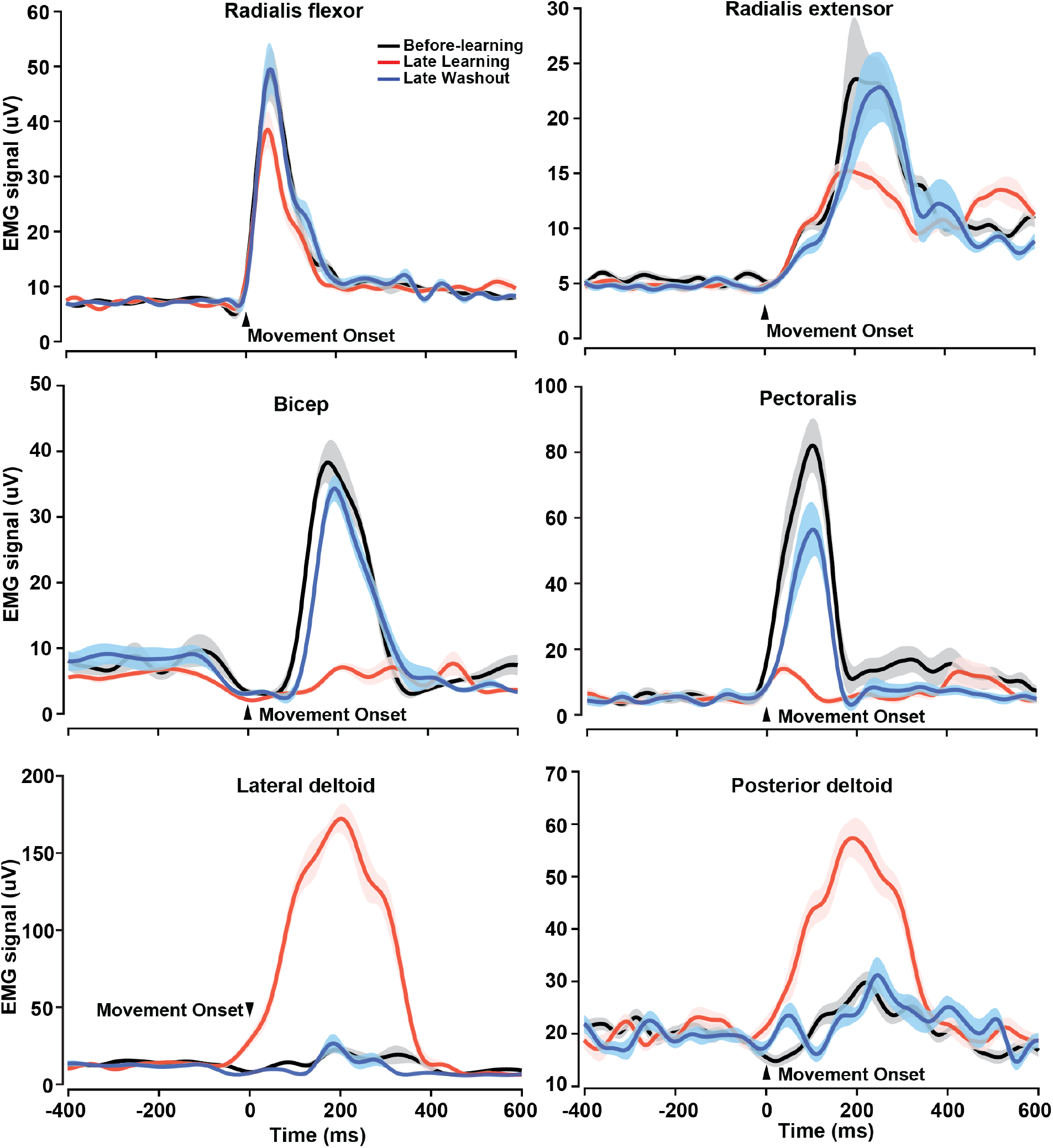
EMG signals of 6 upper limb muscles (bicep, radialis flexor, radialis extensor, pectoralis, posterior deltoid, lateral deltoid) in before-learning, learning, and washout blocks. Time zero, movement onset. One example condition: CW curl field applied to down reaches. EMG signals do not show signs of muscle co-contraction in late-learning trials (red). Muscle activity during the preparatory period remains flat and around the same level across all blocks *(P* > 0.3 for all pairs of comparison except for *P* < 0.0001 when comparing late-learning bicep activity with before-learning/late-washout bicep activity). Muscle activity patterns during before-learning (black) and late-washout trials (blue) are very similar. Shaded area, s.e.m across trials. EMG activity shows similar temporal patterns to previous intramuscular recordings^53,61^.

**Supplementary Figure 4.**
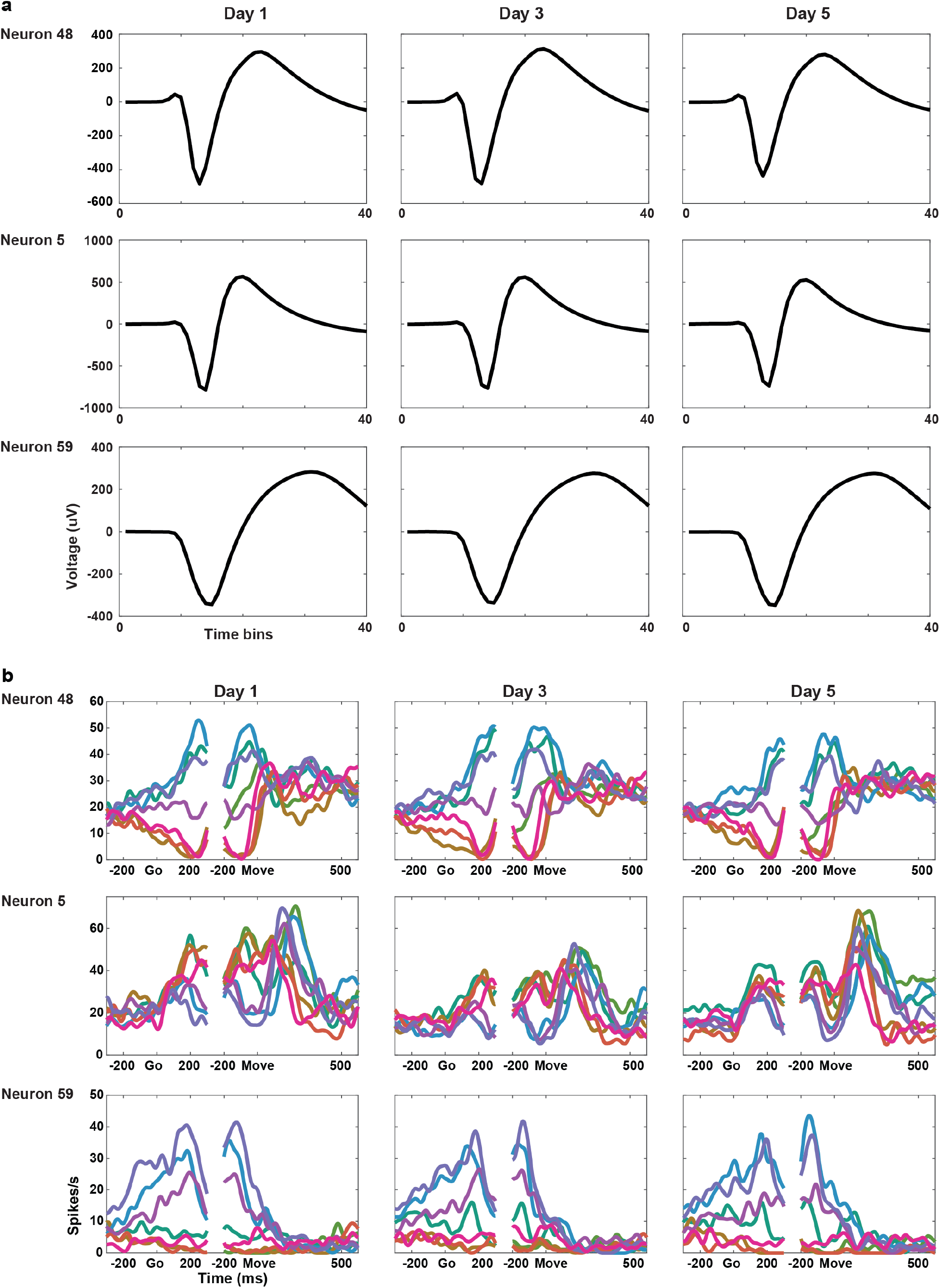
Stability of example spike waveforms and PSTHs over five successive sessions. The same 71 units from Monkey U Utah-array recordings were selected post-hoc by comparing waveform correlations and tracked over five sessions. **a**, All selected units have nearly-identical waveforms. Waveforms of three representative units are shown. **b**, Most selected units have similar direction-tuning properties for before-learning reaches across sessions. PSTHs of three representative units for eight reach directions are shown. Go, go cue. Move, movement onset.

**Supplementary Figure 5.**
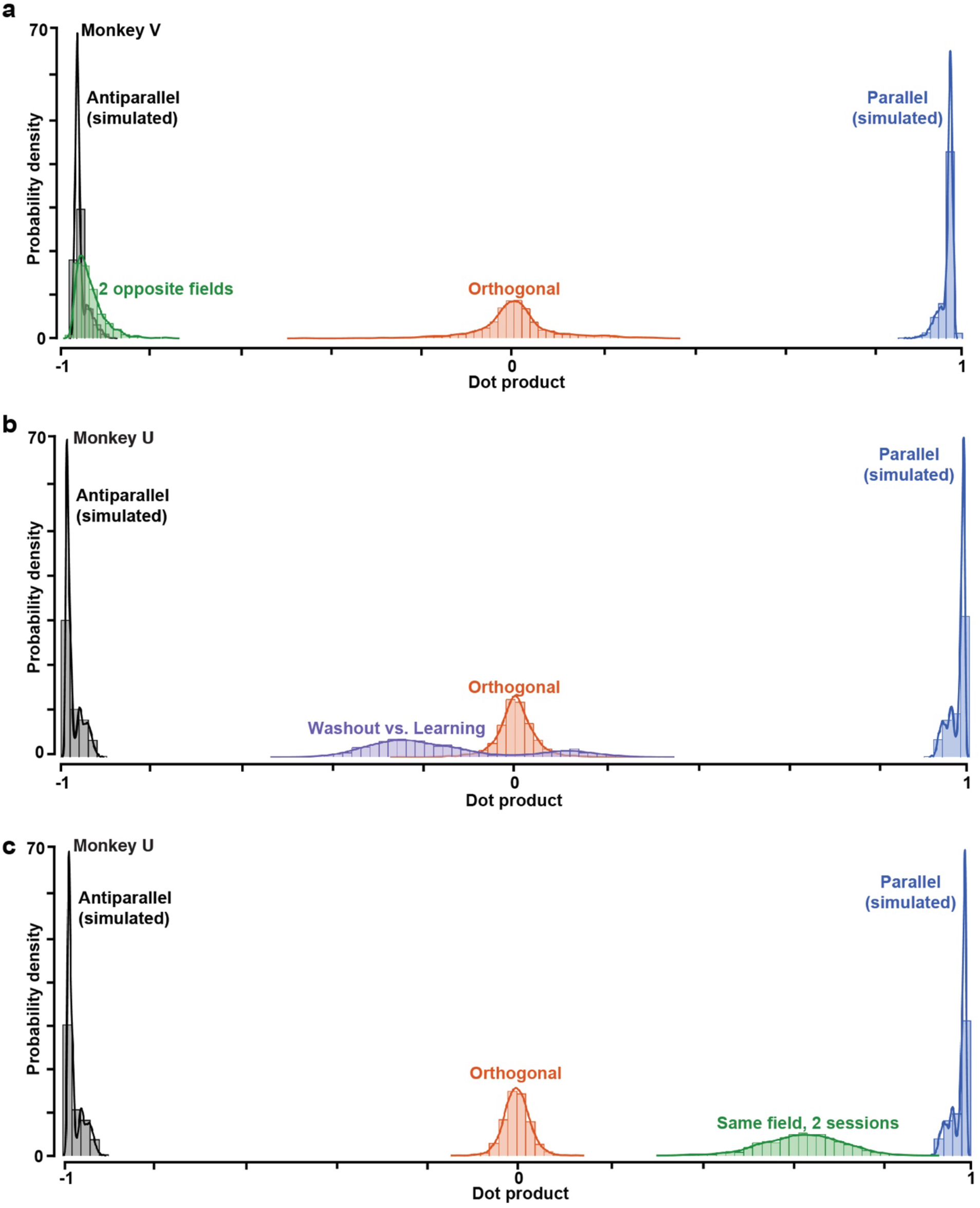
Distributions of dot products between uniform shifts. **a-c**, Distributions of dot products between uniform-shift axes when **a**, learning two opposite fields for one reach target (Monkey V, green, around −1), **b**, learning and washing out one curl field (Monkey U, purple, close to 0), and **c**, learning the same curl field in two sessions 18 days apart (Monkey U, green, close to 1). We compare them with hypothetical distributions of dot products between uniform shifts predicted by orthogonal (red), parallel (blue), and antiparallel (black) relationships simulated with real data (see Methods).

**Supplementary Figure 6.**
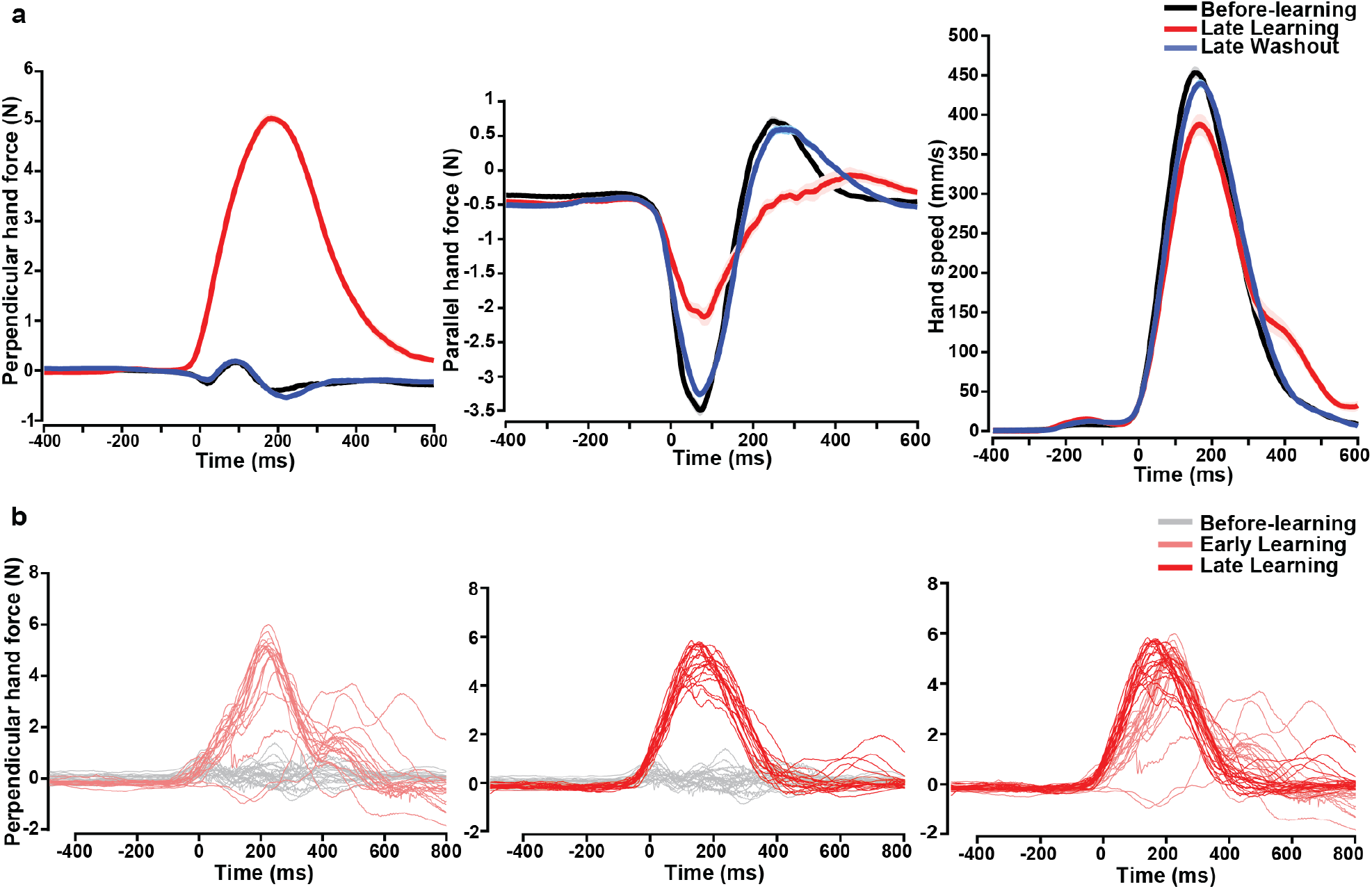
Single-trial (upper panel) and trial-averaged (lower panel) hand forces in different blocks from one representative session. **a**, Hand forces perpendicular to (left) and parallel with (middle) the reach direction change greatly in late-learning trials (red curve) but revert to before-learning level (blue curve) in late-washout trials (black curve). Right, peak hand speed slightly decreases in learning trials but reverts to before-learning level in late washout trials. Shaded area, s.e.m. across trials. **b**, Compensatory hand forces perpendicular to the reach direction increase from the very beginning of the learning block (light red curves, first 20 trials during learning) compared to the before-learning trials (gray curves), showing the immediate online feedback control to correct the perturbed movement. Hand forces in late-learning trials (dark red curves, last 20 trials during learning) show a more stereotypical, less variable temporal pattern with an earlier onset than in early-learning trials^61^. **a, b**, Time zero, movement onset.

**Supplementary Figure 7.**
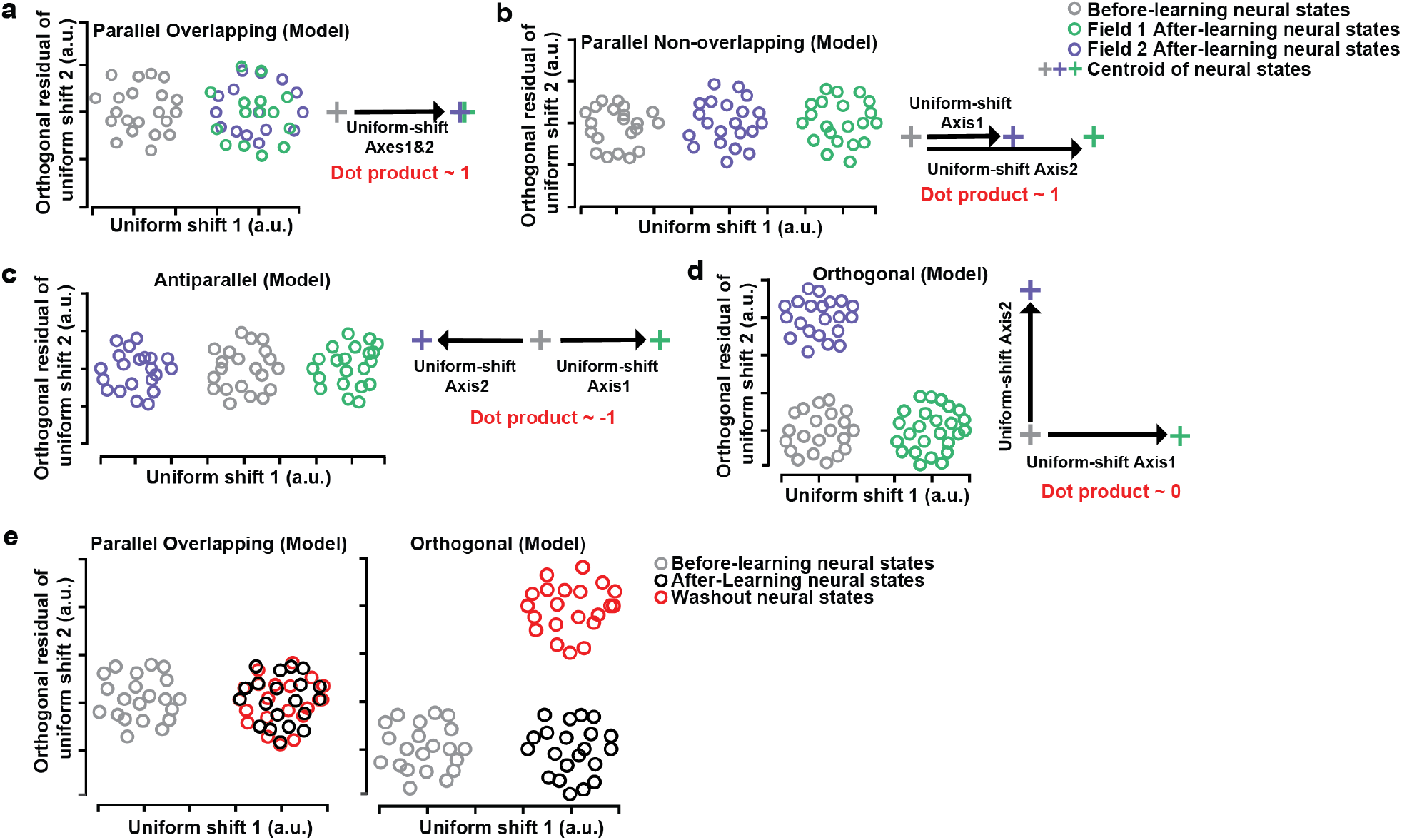
Hypothetical geometric relationships of uniform shifts for learning different curl fields or for washout. **a-d**, Hypothetical models illustrating geometric relationships of uniform shifts for learning different curl fields. Left panels, illustrations of before-learning and after-learning neural states when learning two distinct curl fields. Right panels, the definition of uniform-shift axes in each case. **a**, Parallel overlapping: the uniform shifts of neural states for both fields are in the same direction, and two after-learning neural repertoires are mixed. **b**, Parallel non-overlapping: the uniform shifts of neural states for both fields are in the same direction, and two after-learning neural repertoires are separated. **c**, Antiparallel: the uniform shift of neural states for learning one field is opposite to that for learning another field. **d**, Orthogonal: uniform shifts for two fields in directions that are independent. **e**, Hypothetical models illustrating geometric relationships of uniform shifts for learning and washout of one curl field. Parallel overlapping (left): washout neural states (red) do not move further from after-learning states (black). Orthogonal (right): washout neural states (red) move away from after-learning states (black) along another axis orthogonal to the learning uniform-shift axis.

**Supplementary Figure 8.**
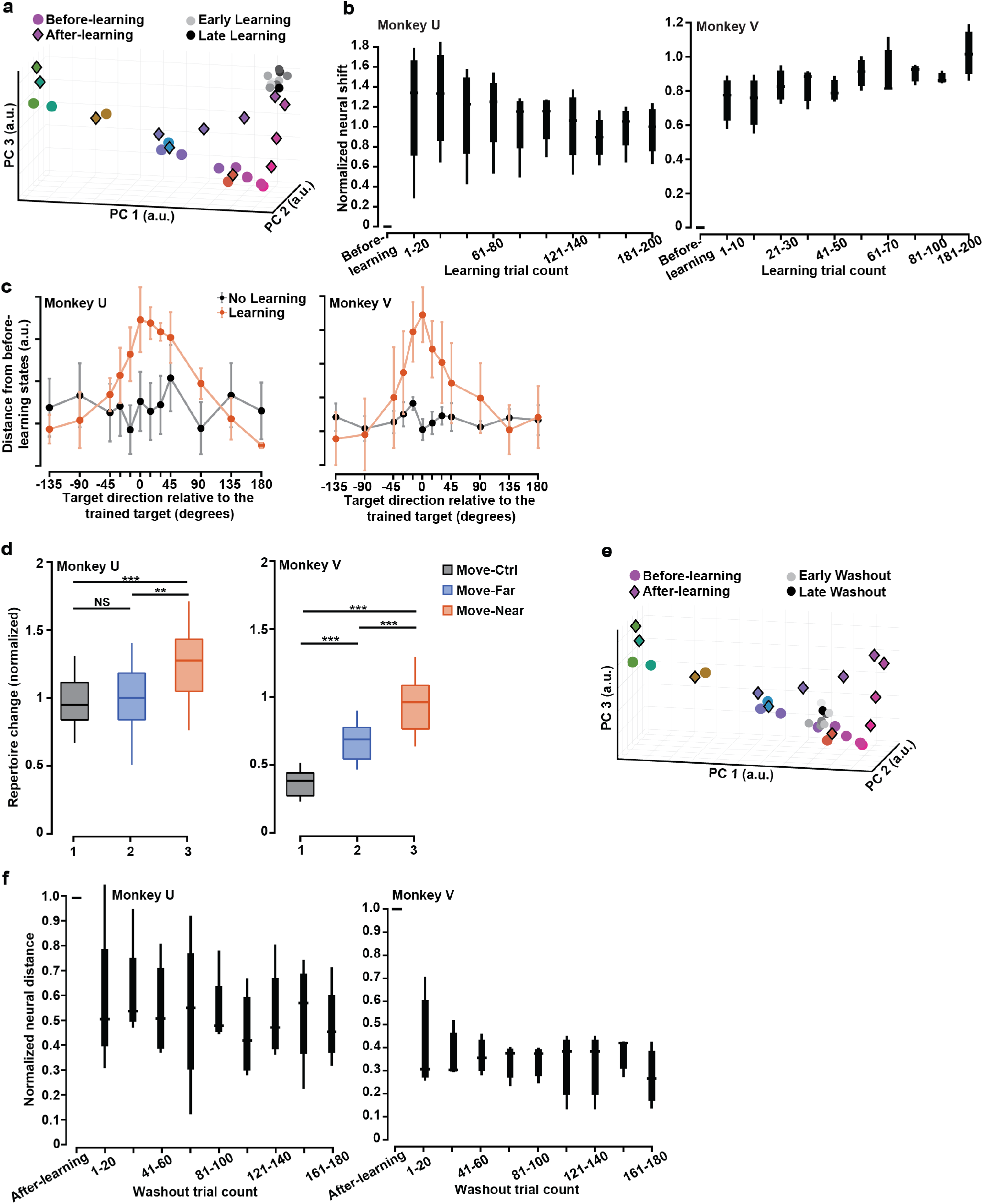
Peri-movement neural states projected to PCs 1-3 in before-learning, learning and washout blocks, their local generalization of learning and non-uniform repertoire change. **a**, Peri-movement states for before-learning (color circles), learning (grey circles), and after-learning error-clamp reaches (color diamonds) projected to PCs 1-3. After-learning neural states leave the before-learning repertoire, and this shift is local for reach targets that are close to the trained target. **b**, Quantification of neural state shift during learning along the axis that connects before-learning and after-learning states of the trained target, normalized against the distance between these two states (averaged every 10 or 20 trials. Cuzick’s test: Monkey U, *P* = 0.032; Monkey V, *P* = 3.92 x 10^-5^). **c**, The local shift of peri-movement neural states, quantified as the Euclidian distance between before-learning and after-learning states of each reach target, shows bell-shaped generalization pattern. Error bars, s.e.m. across sessions. **d**, Peri-movement activity patterns show a significantly greater repertoire change for the trained target and near targets within 45 degrees from the trained target (red, Move-Near) than far targets more than 45 degrees from the trained target (blue, Move-Far). Black (Move-Ctrl), the repertoire change value in control sessions when monkeys did thousands of center-out reaches without any force field. One-sided Wilcoxon rank-sum test: Monkey U, *P_12_* = 0.26, *P_13_* = 4.52 x 10^-6^, *P_23_* = 0.002; Monkey V, *P_12_* = 3.70 x 10^-7^, *P_13_* = 6.02 x 10^-9^, *P_23_* = 5.29 x 10^-4^. **e**, Peri-movement neural states in the same PCA subspace for early (lighter grey) and late (darker grey) washout trials. **f**, During washout, the distance between washout and before-learning states decreases significantly along the axis that connects before-learning and after-learning states of the trained target. All distances are normalized against the distance between before-learning and after-learning states of the trained target (averaged every 20 trials. Cuzick’s test: Monkey U, *P* = 0.0077; Monkey V, *P* = 0.0028).

